# A Genome-Wide Association Analysis Reveals a Role for Recombination in the Evolution of Antimicrobial Resistance in *Burkholderia multivorans*

**DOI:** 10.1101/313205

**Authors:** Julio Diaz Caballero, Shawn T. Clark, Pauline W. Wang, Sylva L. Donaldson, Bryan Coburn, D. Elizabeth Tullis, Yvonne C.W. Yau, Valerie J. Waters, David M. Hwang, David S. Guttman

## Abstract

Cystic fibrosis (CF) lung infections caused by members of the *Burkholderia cepacia* complex, such as *Burkholderia multivorans*, are associated with high rates of mortality and morbidity. We performed a population genomic study of 111 *B. multivorans* sputum isolates from a single CF patient through three stages of infection including the initial incident infection, deep sampling of a one-year period of chronic infection, and deep sampling of a post-transplant recolonization. We reconstructed the evolutionary history of the population and used a lineage-controlled genome-wide association study (GWAS) approach to identify genetic variants associated with antibiotic resistance. We found that the incident isolate was more susceptible to agents from three antimicrobial classes (β-lactams, aminoglycosides, quinolones), while the chronic isolates diversified into distinct genetic lineages with reduced antimicrobial susceptibility to the same agents. The post-transplant reinfection isolates displayed genetic and phenotypic signatures that were distinct from sputum isolates from all CF lung specimens. There were numerous examples of parallel pathoadaptation, in which individual loci, or even the same codon, were independently mutated multiple times. This set of loci was enriched for functions associated with virulence and resistance. Our GWAS approach identified one variant in the *ampD* locus (which was independently mutated four times in our dataset) associated with resistance to β-lactams, and two non-synonymous polymorphisms associated with resistance to both aminoglycosides and quinolones, affecting an *araC* family transcriptional regulator, which was independently mutated three times, and an outer member porin, which was independently mutated twice. We also performed recombination analysis and identified a minimum of 14 recombination events. Parallel pathoadaptive loci and polymorphisms associated with β-lactam resistance were over-represented in these recombinogenic regions. This study illustrates the power of deep, longitudinal sampling coupled with evolutionary and lineage-corrected GWAS analyses to reveal how pathogens adapt to their hosts.

## Author Summary

Cystic fibrosis (CF) is a common lethal genetic disorder that affects individuals of European descent and predisposes them to chronic lung infections. Among the organisms involved in these infections, bacteria from the *Burkholderia cepacia* complex (BCC) are often associated with poor clinical prognosis. This study examines how the most prevalent BCC species among CF patients, *B. multivorans*, evolves within a CF patient and identifies mutations underlying antibiotic resistance and adaptation to both the native CF lung and a non-CF lung allograft. We demonstrate that *B. multivorans* can diversify phenotypically and genetically within the CF lung, with a complex population structure underlying a chronic infection We noted that isolates collected after the patient was re-infected post-transplant were more closely related to descendants of the incident clone than to those recovered in the weeks prior to transplant. We used a genome-wide association method to identify genes associated with resistance to the β-lactam antibiotics: aztreonam and ceftazidime. Many of these variants were found in regions that show patterns of recombination (genetic exchange) between strains. We also found that genes which were mutated multiple times during overall infection were more likely to be found in regions showing signals consistent with recombination. The presence of multiple independent mutations in a gene is a very strong signal that the gene helps bacteria adapt to their environment. Overall, this study provides insight into how pathogens adapt to the host during long-term infections, specific genes associated with antibiotic resistance, and the origin of new and recurrent infections.

## Introduction

The *Burkholderia cepacia* complex (BCC) describes a highly diverse group of at least 20 closely related species within the genus *Burkholderia* that can cause serious opportunistic infections in humans [1, 2]. Individuals with the fatal genetic disease cystic fibrosis (CF) are particularly susceptible to chronic BCC infections, which are commonly associated with rapid decline in lung function, high rates of mortality and poor post-transplant outcome [3, 4]. Of the BCC species, *Burkholderia multivorans* and *Burkholderia cenocepacia* account for 85-97% of all BCC found in CF patients [5]; however, *B. multivorans* infections have surpassed *B. cenocepacia* in prevalence over the past decade [6]. Many BCC that are CF-associated are intrinsically virulent and antibiotic resistant and require strict infection control practices, as they can be transmitted between patients [7-10]. Despite a wealth of knowledge describing the molecular basis of these pathogenic properties and their evolution in strains of the well-studied *B. cenocepacia*, little is known about the factors that govern these attributes in *B. multivorans* [9].

Dissecting the molecular basis of complex adaptive traits in bacterial pathogens, such as antimicrobial resistance, can be difficult as a single phenotype may be influenced by a large number of loci that interact with each other as well as their environment. Resistance in the BCC is associated with alterations to outer membrane permeability, the expression of multidrug efflux pumps and β-lactamases, and diversification of antimicrobial targets [11]. Consequently, methods that focus on identifying polymorphisms in single genes with large effects may miss the majority of loci that modulate phenotypes in more subtle ways. The development of genome-wide association studies (GWAS) has expanded our ability to identify loci of small effect size that have been associated with numerous diseases and other related phenotypes of interest in humans [12, 13]. In contrast, the application of GWAS to analyze bacterial behaviors has been slower to gain traction for a number of inter-related reasons: 1) clonal reproduction of microbes leads to confounding associations due to common ancestry, often referred to as population structure; 2) recombination in bacteria, which is more analogous to gene conversion than eukaryotic recombination, occurs at variable rates among different species and is not linked to reproduction; 3) the unpredictable nature of recombination results in the erratic breakdown of linkage disequilibrium between selected sites and distal neutral sites; and 4) selection can be extremely strong, resulting in the relatively rapid fixation of not only a selected allele, but entire genomes due to the linkage disequilibrium [14, 15].

Nevertheless, several recent studies have proposed novel approaches to overcome these challenges. These methods include using cluster membership [16-18], phylogenetic history [15, 19, 20], or lineage effects [21] to differentiate mutations leading to a phenotypic outcome from mutations related to the genetic background of the bacterial population. While these methods hold tremendous promise for identifying genetic variation underlying bacterial phenotypes of interest, they generally focus on cross sectional sampling of diverse isolates and populations. Their power has not been established for the fine-scale analysis of individual bacterial populations evolving over short time scales, with strong positive selection and restricted recombination [14, 22]. The application of fine-scale evolutionary analysis to bacterial populations is especially important in the context of clinically significant pathogen infections, where evolution is associated with adaptation to the host environment and antimicrobial treatment [23].

In this study, we take a fine-scale approach to microbial GWAS to examine the genetic basis of antimicrobial resistance within a *B. multivorans* population that had been sampled longitudinally from a single patient over a ten-year period. We characterized the genomic diversity in this population and assessed associations between all genetic variants and multiple antibiotic resistance phenotypes. Using a clustering-based approach to control for population structure and linkage disequilibrium, our analysis identified single nucleotide polymorphisms (SNPs) that were associated with resistance to β-lactams, aminoglycosides, and quinolones. In addition, we found that both multiply-mutated loci (those that are targets of parallel pathoadaptation) and β-lactam resistance-associated variants were overrepresented in recombinogenic regions of the *B. multivorans* genome

## Results

For our evolutionary analysis and GWAS, we used a series of *B. multivorans* isolates that were cultured from respiratory specimens obtained from an adult male with CF (CF170, being followed by the CF Clinic at St. Michael’s Hospital, Toronto, Canada). In a ten-year period, patient CF170 acquired an incident (i.e. initial) lung *B. multivorans* infection, developed a chronic *B. multivorans* lung infection, received a double lung transplant, and finally experienced a *B. multivorans* re-colonization of the allograft three years post-transplant. Isolates from each of these three phases of his *B. multivorans* infection are represented in this study (Fig 1). We defined these isolates as 1) the single isolate recovered from the patient’s first infection – the ‘incident infection’ isolate; 2) 100 isolates collected six to seven years post-incident infection from ten sputum specimens (ten isolates per specimen) over approximately a one-year period – the ‘chronic infection’ isolates; and 3) ten isolates collected from a single expectorated sputum sample ten years after the incident infection, and three years after the patient underwent a double lung transplant – the ‘post-transplant’ isolates. Patient CF170 was being treated with alternating cycles of antibiotic therapy while chronically infected, with 13 antibiotics being administered at different intervals and durations over the course of the chronic infection sampling period (Fig 1). The genomes of all 111 isolates were whole-genome sequenced on the Illumina platform, yielding a median coverage depth of 117X (S1 Fig). Multi-locus sequence typing was performed *in silico* by extracting seven loci from the whole genome sequence data *(atpD, gltB, gyrB, recA, lepA, phaC, trpB)* and comparing them to the *Burkholderia cepacia* complex MLST Databases in pubMLST. This analysis revealed that all isolates were clonally related and of the sequence type ST-783 [24].

**Fig 1.**
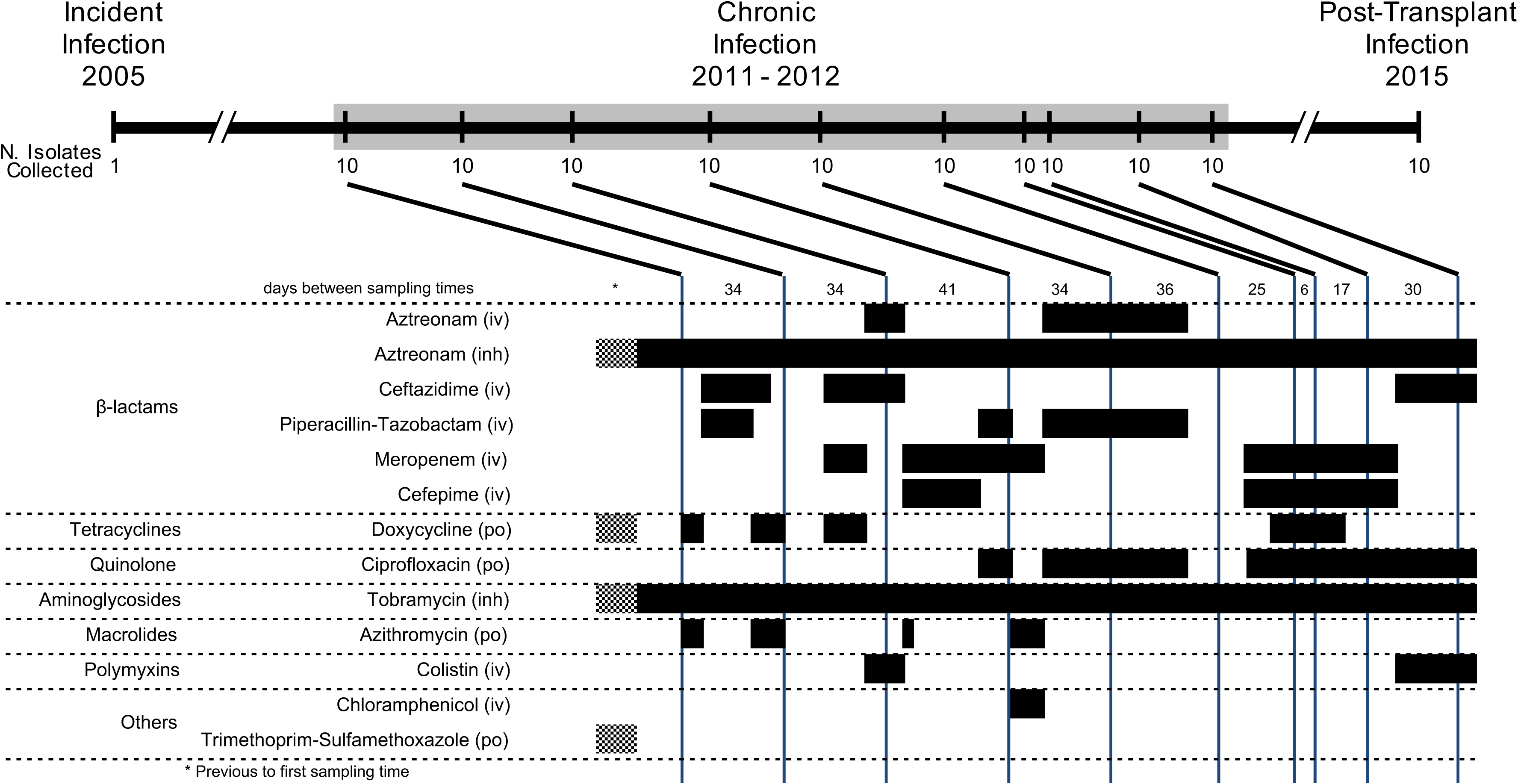
Time course of *B. multivorans* infection in study patient CF170. A total of 111 *B. multivorans* isolates from twelve collection times were used in this study (1 isolate from the initial infection, 10 isolates from each of 10 sputum samples collected during chronic infection, and 10 isolates from a sputum sample obtained during a post-transplant infection). Antibiotic treatment history during the chronic infection period is shown in the lower panel. Black bars indicate antibiotic administration, while hashed bars indicate intermittent exposure in that time block (only relevant prior to the start of chronic sampling). The method of antibiotic administration is shown as intravenous (iv), inhaled (inh), or oral (po).

### Genomic diversity and phylogenetic analysis suggest underlying population structure

The *de novo* genome assembly of a single isolate recovered from the third chronic infection sputum sample was used as the reference for the mapping assembly of all other isolates. This particular isolate was chosen as the reference since it had the best overall *de novo* assembly metrics. The reference assembly consisted of 6,444,123 bases across 26 contigs, which were pseudo-scaffolded against the complete genome of *B. multivorans* ATCC 17616 (Fig 2A). Through a conservative variant calling pipeline [25], a total of 1,892 SNPs and 328 indels segregating among the 111 isolates were identified, with 1,039, 672, and 180 SNPs being found on chromosomes, 1, 2, and 3 respectively. Only a single SNP was found in a contig which did not map to the ATCC 17616 genome. Overall, 740 (39.1%) SNPs and 163 (49.9%) indels were parsimonious informative (PI, i.e. non-singleton), and 226 (11.9%) SNPs and 99 (30.2%) indels segregated in at least two sampling time points. From the 1,892 SNPs, 70.5%, 15.6%, and 13.9% were non-synonymous, synonymous, and intragenic substitutions respectively (Fig 2C). 52.1% of the intergenic SNPs were found in putative regulatory regions (defined as the intergenic region within 150 bases from the start codon of any gene). The population showed a genetic diversity average of 123.62 ± 120.98 (number of SNP differences, mean ± standard deviation) pairwise differences. The distribution of these difference suggested an underlying population structure since genetic diversity was not uniform even among isolates from the same specimen (S2 Fig).

**Fig 2.**
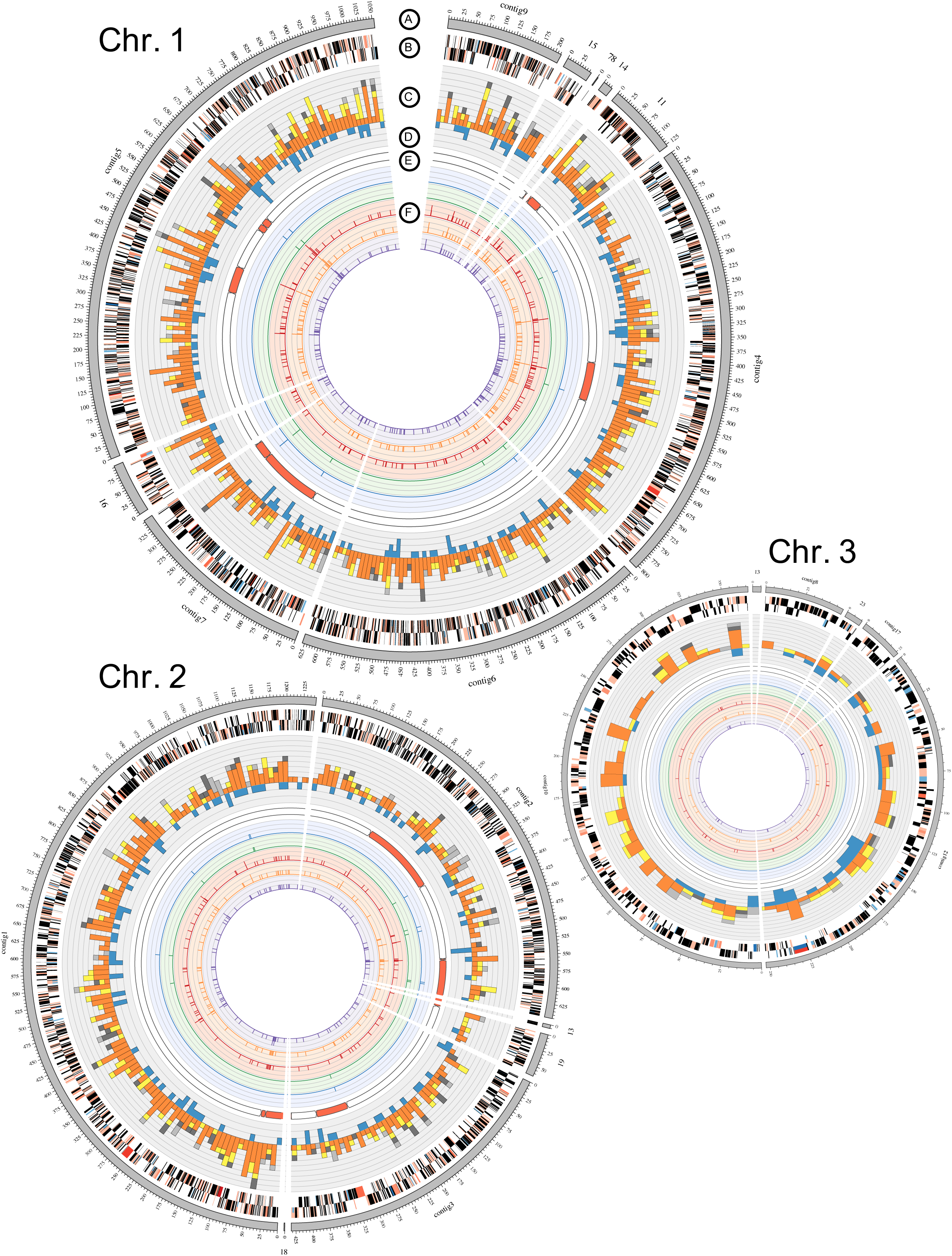
Genomic Characterization of 111 *B multivorans* isolates. (A) Contigs (gray outer ring) of the *de novo* reference were arranged according to the three chromosomes of the complete genome of *B. multivorans* ATCC 17616. This genome was obtained from expectorated sputum collected in the third chronic infection sample. (B) Genome annotation according to RAST. (C) SNP count per 10 Kb as a function of their location in the contigs. Non-synonymous (orange), synonymous (yellow), putative regulatory (dark grey) and intergenic (light grey). (D) Indel (blue) count per 10 Kb. (E) Recombinogenic regions, as predicted by DnaSP Hudson-Kaplin four gamete test, are shown as red blocks. (F) Variants Associated with Antibiotic Resistance. From outermost to innermost ring: aztreonam and ceftazidime (β-lactam), amikacin and tobramycin (aminoglycoside), and ciprofloxacin (quinolone). This figure was prepared with circus v. 0.69 [69].

We reconstructed the core genome phylogenetic relationships among all isolates using an alignment of the 1,892 SNPs and a Bayesian approach (Fig 3A). The root of this tree was identified by adding *B. multivorans* ATCC17616 to the analysis. The tree topology indicates that the incident infection isolate diverged from the other 110 isolates at the base of the tree. The ten isolates from the post-transplant sample are again highly divergent (relative to the total diversity) and form a basally branching, monophyletic clade, while the chronic sample isolates form a less divergent, monophyletic clade. Moreover, there seem to be subgroups among the chronic infection isolates suggesting population structure. This structure is also observed in a network-based phylogenetic approach (S3 Fig), where two groups of isolates from the chronic infection sampling cluster in a star-like phylogeny. Star phylogenies are characterized by roughly equal divergence from the common ancestor, and are associated with recent purges in genetic variation [26].

**Fig 3.**
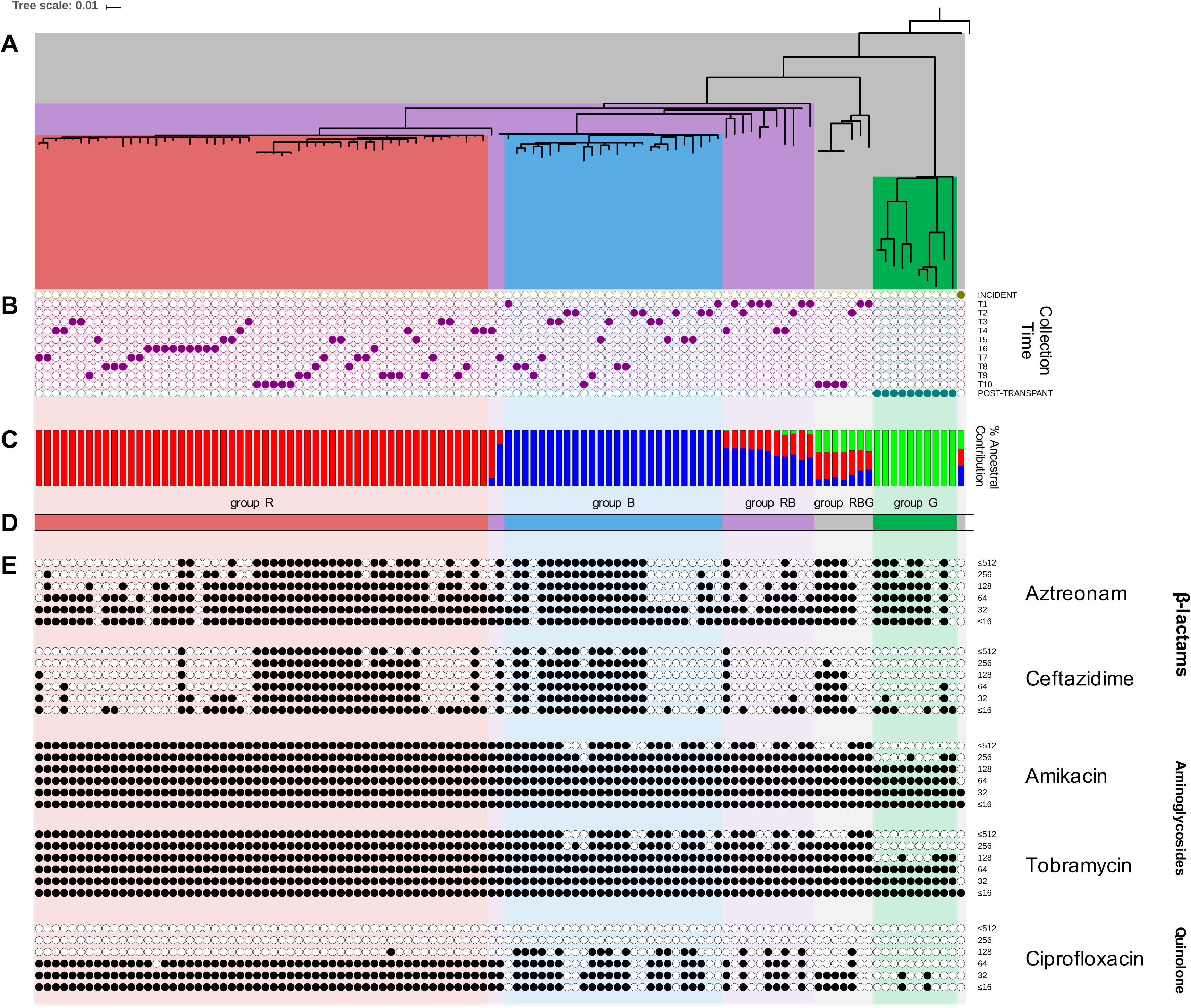
Population structure and antibiotic resistance profiles. (A) Phylogenetic relationships of the 111 *B. multivorans* isolates were estimated employing a Bayesian approach based on genome-wide single nucleotide polymorphisms (SNPs). (B) Time of collection for each isolate. (C) Population structure analysis as assessed by Structure v2.3.4 with three expected ancestral subpopulations. Ancestral subpopulations are coded as red (R), blue (B), and green (G). (D) Isolates are grouped based on their ancestral composition. Group R, B, G, RB, and RBG are shaded in red, blue, green, purple, and grey respectively. (E) Antibiotic susceptibility for each isolate, the highest black circle represents the MIC μg/mL), to the β-lactams: aztreonam and ceftazidime, the aminoglycosides: amikacin and tobramycin, and the quinolone: ciprofloxacin are shown as filled circles at six different concentration thresholds. This figure was elaborated at the interactive tree of life (iTOL) website v. 3 [70].

### Population structure analysis clusters the isolates into five groups

We used the Monte Carlo Markov Chain analysis of SNPs and indels implemented in STRUCTURE to infer population structure among the 111 isolates. We identified the lowest number of subpopulations that maximized the likelihood of data; hence determining the underlying population structure in the data without overestimating the number of subpopulations [27]. There were three subpopulations that arose from single common ancestors, which we labelled groups R, B, and G, comprising 54, 26, and 10 isolates, respectively (Fig 3C-D). The ancestral composition of the incident isolate and seven of the chronic infection isolates, recovered at collection points T1, T2 and T10, resembled a combination of the three identified subpopulations. This group of isolates was labeled RBG. Another group labeled RB (13 isolates) has an admixed ancestry from the ancestral subpopulations of R and B.

Isolates from groups RBG and RB were found in low frequencies through different samples from the chronic infection period (Fig 3B). In contrast, isolates from group R or B were more dominant in this same period. The isolates from group R were first observed at the third time point of the chronic infection samples, and they remained the most abundant group in subsequent chronic samples (Fig 4). In contrast, the abundance of group B isolates decreased over time. The genetic diversity, measured as number of SNPs, significantly differed between these groups (one-way ANOVA: F(4,1902) = 1,426.133, p-value < 0.0001), with group G (those recovered exclusively post-transplant) being the most diverse, followed by groups RBG and RB, then groups R and B (S4a Fig).

**Fig 4.**
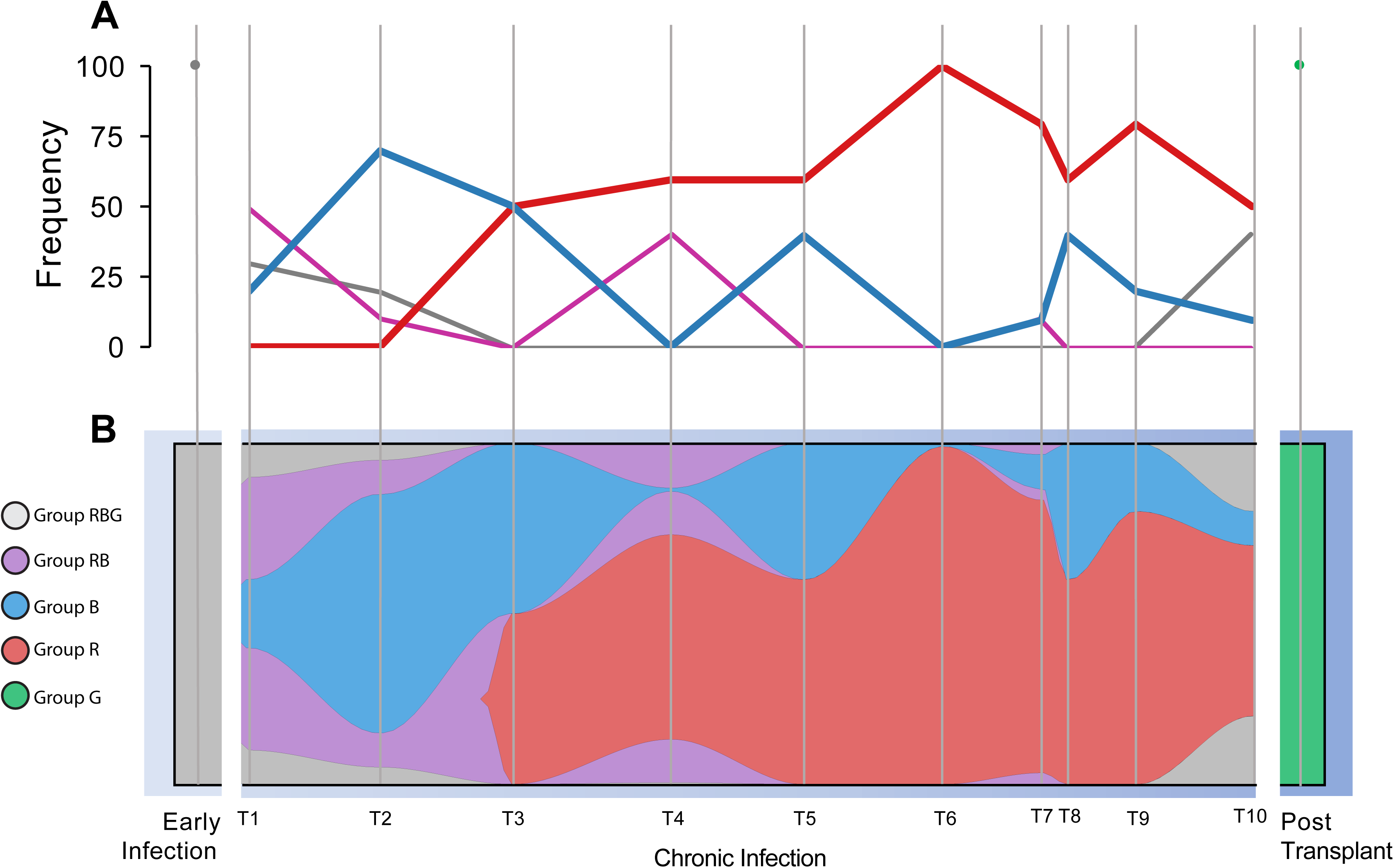
Population genomics of the community over time. Groups R, B, G, RB, and RBG are coloured in red, blue, green, purple, and grey respectively. (A) Frequency of each group over time. (B) The clonal graph was created with the assumption that RGB is the group of isolates resembling the ancestor of all the isolates, and RB is the group of isolates resembling the ancestor of group R and B. The distance between sample times is relative to the actual number of days between them. This plot was created using fishplot v. 0.3 [71].

The time to the most recent common ancestor (tMRCA) calculated as days before the last sample for all isolates and the various STRUCTURE-defined groups is shown in Supplementary Figure 4c. This analysis shows that the RGB group, which includes all of the chronic infection isolates as well as the post-transplant isolates, coalesced to a common ancestor at roughly the same time as the full isolate collection, including the incident infection (S4c Fig). This result supports the hypothesis that the infection of the transplanted lung came from the same source as the original incident isolate, despite being separated by approximately ten years, as opposed to a clone that persisted and diversified in the lung of the patient during chronic colonization. Additionally, it appears that groups R and B diverged at approximately the same time (S4c Fig). Unfortunately, we are unable to determine if these were allopatric populations that colonized distinct regions in the lung, or sympatric populations that coexisted within the same compartment due to our sampling of expectorated sputum.

### Selection analysis supports positive selection in the population

We determined the ratio of non-synonymous to synonymous substitutions (d_N_/d_S_) as an estimate of selection. Since we expect that time has allowed natural selection or genetic drift to have acted on the multi-time segregating mutations more so than on variants that segregate in a single sample, we determined the d_N_/d_S_ both for all SNPs in each group, as well as for only those that segregate in at least two time-points – ‘multi-time’ SNPs (S4b Fig). The d_N_/d_S_ for the overall population was 1.35 (95% confidence interval, CI = 1.19-1.53) and 1.34 for multi-time SNPs (CI = 0.94-1.96), which may indicate weak positive selection, or simply the segregation of mildly deleterious variants. Only groups R and RB multi-time SNPs showed d_N_/d_S_ above the neutral expectation of 1.0 (group R d_N_/d_S_ = 2.05, CI = 0.57-11.15, group RB d_N_/d_S_ = 2.38, CI = 1.08-6.18), although the confidence intervals for the group R are quite large. All other groups had d_N_/d_S_ ratios only slightly elevated (ranging from 1.04-1.63), although the differences between groups were not statistically significant.

Further support for positive selection comes from a significantly negative Tajima’s D test (D = − 2.21, P < 0.01) and Fu and Li’s tests (D* = −6.11, P < 0.02; F* = −5.20, P < 0.02). While all three of these results can be explained by both positive selection and recent population expansion, the combination of these results with the high nucleotide diversity and d_N_/d_S_ > 1.0 is most consistent with positive selection.

### GWAS identification of variants associated with antibiotic resistance

We assumed that the intensive antibiotic exposure during the chronic infection sampling period would result in strong selection for resistance-associated genotypes in *B. multivorans*. Minimum inhibitory concentrations (MICs) for two β-lactams (aztreonam, ceftazidime), two aminoglycosides (tobramycin and amikacin), and the fluoroquinolone ciprofloxacin were determined for all isolates. Isolates from the three phases of infection had distinct susceptibility profiles. The incident isolate had MICs of 8 μg/mL or less for all agents tested, while all chronic infection and post-transplant isolates had higher MICs for both aminoglycosides (t-test p < 0.0001, Fig 3E), but variable MICs for β-lactams and fluoroquinolone tested (range: ≤8 to >512 μg/mL).

The 1,892 SNP positions segregating among the 111 isolates were grouped in 150 distinct mutational profiles (i.e. one or more SNP positions that share the same pattern of reference vs. alternative base among the strain collection, S5 Fig). Prior to population control, each of these mutational profiles was examined for a statistical association to the five tested antibiotics at six different levels of resistance and these associations were corrected for multiple testing by taking into consideration the number of tests. Five mutational profiles (comprising 17 SNPs) associated with resistance to both β-lactam antibiotics, and one mutational profile (comprising 2 SNPs) associated specifically with ceftazidime (S6 and S7 Fig). Ten mutational profiles (comprising 250 SNPs) were associated with resistance to amikacin, tobramycin, and ciprofloxacin. Additionally, two mutational profiles (comprising 31 SNPs) associated with resistance to both aminoglycosides, and four mutational profiles (comprising 33 SNPs) associated specifically with ceftazidime.

Next, we tested these variants against population structure controls, counting only those associated variants that were observed in multiple subpopulation groups as determined by the population structure analysis. This criterion could be satisfied by one of two mechanisms: 1) the mutations arose in the subpopulations through multiple independent mutational events, or 2) they arose in a common ancestor of multiple subpopulations and have been maintained in multiple lineages while being lost in others. Out of all mutational profiles associated with elevated MICs for both β-lactams, one (comprising a single SNP) passed the population structure control (S6b Fig). This SNP was found in 20.4% of isolates in group R, and 50% of isolates in group RBG. This variant leads to a non-synonymous amino acid substitution in the sequence of the *ampD* gene (BMUL_2790), a locus extensively studied for its role in resistance to β-lactams [28, 29]. This mutation was predicted to have a deleterious effect on AmpD by PROVEAN analysis (score = −8.0, S8a Fig). In fact, the *ampD* locus was independently mutated four other times within our dataset. A second SNP in *ampD* was found in a mutational profile that was similarly associated with β-lactam MICs; nevertheless, it failed to pass the population structure control. Additionally, two mutational profiles associated to the aminoglycosides and ceftazidime showed evidence of multiple independent polymorphic events (S6e Fig). One of these mutational profiles, which comprises a single SNP, is represented by a non-synonymous substitution in an *araC* family transcriptional regulator locus (BMUL_3951). PROVEAN analysis indicates that this mutation is unlikely to have a deleterious effect on the locus (score = 6.906). The second mutational profile, again including only a single SNP, gave rise to a non-synonymous substitution in locus BMUL_3342, which is annotated as an outer member protein (porin). While this mutation is not expected to end in a deleterious effect (PROVEAN score = 3.273), it occurs in a locus that is independently mutated two other times.

### Additional variants associated with pathoadaptation can be detected by identifying multi-mutated loci

Loci that are independently mutated multiple times provide strong evidence of selection by parallel pathoadaptation [30]. We observed 328 loci that were independently mutated multiple times in our collection (Table 1). Given the genome size and the total number of polymorphisms (both SNPs and indels), we only consider the 62 loci with three or more independent mutations to be statistically significant (p-value < 0.05/[1,892 SNPs + 328 indels = 2220 polymorphisms]). 184 SNPs (9.7%) and 26 indels (7.9 %) were found in these 62 loci. We excluded the possibility that multiply mutated loci showed excess polymorphism simply due to an increased mutational rate by examining the mutational class spectrum for the multiply mutated loci relative to the genome-wide average. While the rate of non-synonymous, synonymous and intergenic mutations among all 1,892 SNPs is 70.5%, 15.6%, and 13.9% respectively, the mutational class spectrum of the SNPs found among multiply mutated loci is 83.1% non-synonymous, 11.7% synonymous, and 3.2% intergenic substitutions. Therefore, the mutational class distribution of SNPs found in multiply mutated loci is significantly skewed toward an excess of non-synonymous mutations (P < 0.0001, chi-square test).

**Table 1.** Parallel Pathoadapted Loci with Multiple Independent Mutations.

| Locus | Encoded Protein | No. of SNPs/Indels | Probability <sup>a</sup> |
| --- | --- | --- | --- |
| BMUL_0641 | Probable transcription regulator protein of MDR efflux pump cluster | 7/0 | 1.65 X 10<sup>-23</sup> |
| BCEN2424_5592 <sup>c</sup> | Glycosyltransferase 36 | 4/2 | 1.03 X 10<sup>-19</sup> |
| BMUL_4010 | NAD-glutamate dehydrogenase | 5/0 | 6.48 X 10<sup>-16</sup> |
| BMUL_0487 | Hypothetical protein | 5/0 | 6.48 X 10<sup>-16</sup> |
| BMUL_4327 | Porin | 3/2 | 6.48 X 10<sup>-16</sup> |
| BMUL_2790 | N-acetyl-anhydromuranmyl-L-alanine amidase (AmpD) | 5/0 | 6.48 X 10<sup>-16</sup> |
| BMUL_1598 | Amino acid adenylation domain-containing protein | 4/0 | 4.06 X 10<sup>-12</sup> |
| BMUL_0353 | YD repeat-containing protein | 3/1 | 4.06 X 10<sup>-12</sup> |
| BMUL_0449 | Preprotein translocase subunit (SecB) | 4/0 | 4.06 X 10<sup>-12</sup> |
| BMUL_2632 | Chaperone protein (DnaJ) | 4/0 | 4.06 X 10<sup>-12</sup> |
| BMUL_4942 | Signal transduction histidine kinase (CheA) | 3/1 | 4.06 X 10<sup>-12</sup> |
| BMUL_2775 | UDP-N-acetylmuramate--L-alanyl-gamma-D-glutamyl-meso-diaminopimelate ligase | 4/0 | 4.06 X 10<sup>-12</sup> |
| BMUL_1444 | Transcription termination factor (Rho) | 4/0 | 4.06 X 10<sup>-12</sup> |
| BMUL_0954 | Glycoside hydrolase 15-like protein | 4/0 | 4.06 X 10<sup>-12</sup> |
| BMUL_4115 | Outer membrane autotransporter | 4/0 | 4.06 X 10<sup>-12</sup> |
| BMUL_0250 | 50S ribosomal protein L4 (RpID) | 3/0 | 2.55 X 10<sup>-8</sup> |
| BMUL_5547 | Conjugation protein (TrbI) | 2/1 | 2.55 X 10<sup>-8</sup> |
| BMUL_2931 | TPR repeat-containing protein | 3/0 | 2.55 X 10<sup>-8</sup> |
| BMUL_3678 | Integral membrane sensor signal transduction histidine kinase | 3/0 | 2.55 X 10<sup>-8</sup> |
| BMUL_3503 | L-serine dehydratase 1 | 3/0 | 2.55 X 10<sup>-8</sup> |
| BMUL_0690 | RND efflux system outer membrane lipoprotein | 2/1 | 2.55 X 10<sup>-8</sup> |
| BMUL_0663 | Alpha/beta hydrolase fold protein | 3/0 | 2.55 X 10<sup>-8</sup> |
| BMUL_0431 | Histidine kinase | 1/2 | 2.55 X 10<sup>-8</sup> |
| BMUL_4510 | Signal transduction histidine kinase (CheA) | 2/1 | 2.55 X 10<sup>-8</sup> |
| BMUL_1970 | Major facilitator transporter | 3/0 | 2.55 X 10<sup>-8</sup> |
| BMUL_2008 | Major facilitator transporter | 2/1 | 2.55 X 10<sup>-8</sup> |
| BMUL_2621 | DNA mismatch repair protein (mutL) | 1/2 | 2.55 X 10<sup>-8</sup> |
| BMUL_4037 | Esterase | 3/0 | 2.55 X 10<sup>-8</sup> |
| BMUL_3977 | Metallophosphoesterase | 2/1 | 2.55 X 10<sup>-8</sup> |
| BMUL_4949 | Aldehyde dehydrogenase | 2/1 | 2.55 X 10<sup>-8</sup> |
| BMUL_3951 | Transcriptional regulator (AraC) | 3/0 | 2.55 X 10<sup>-8</sup> |
| BMUL_6019 | Cytosine/purines uracil thiamine allantoin permease | 2/1 | 2.55 X 10<sup>-8</sup> |
| BMUL_0307 | Amino acid carrier protein | 3/0 | 2.55 X 10<sup>-8</sup> |
| BMUL_5501 | Cytochrome c oxidase subunit I | 3/0 | 2.55 X 10<sup>-8</sup> |
| BMUL_5087 | Short-chain dehydrogenase/reductase SDR | 3/0 | 2.55 X 10<sup>-8</sup> |
| BMUL_4813 | RNA polymerase sigma factor RpoD | 3/0 | 2.55 X 10<sup>-8</sup> |
| BMUL_3197 | Beta-galactosidase | 3/0 | 2.55 X 10<sup>-8</sup> |
| BMUL_3212 | Feruloyl-CoA synthase | 3/0 | 2.55 X 10<sup>-8</sup> |
| BMUL_3315 | PA-phosphatase like phosphoesterase | 1/2 | 2.55 X 10<sup>-8</sup> |
| BMUL_3752 | Peptidoglycan-binding (LysM) | 3/0 | 2.55 X 10<sup>-8</sup> |
| BMUL_3615 | Aldehyde oxidase | 3/0 | 2.55 X 10<sup>-8</sup> |
| BMUL_1686 | Ribonuclease R | 3/0 | 2.55 X 10<sup>-8</sup> |
| BMUL_4615<sup>b</sup> | Amidophosphoribosyltransferase | 3/0 | 2.55 X 10<sup>-8</sup> |
| BMUL_4605 | UTP-glucose-1-phosphate uridylyltransferase | 3/0 | 2.55 X 10<sup>-8</sup> |
| ABD05_14940<sup>d</sup> | Isochorismatase | 3/0 | 2.55 X 10<sup>-8</sup> |
| BMUL_1431 | GAF modulated sigma54 specific transcriptional regulator (Fis) | 2/1 | 2.55 X 10<sup>-8</sup> |
| BMUL_1377 | N-acetyltransferase GCN5 | 3/0 | 2.55 X 10<sup>-8</sup> |
| BMUL_0964 | DNA polymerase III subunit alpha (DnaE) | 3/0 | 2.55 X 10<sup>-8</sup> |
| BMUL_0692 | Carbohydrate kinase FGGY | 2/1 | 2.55 X 10<sup>-8</sup> |
| BMUL_0477 | Error-prone DNA polymerase (DnaE2) | 3/0 | 2.55 X 10<sup>-8</sup> |
| BMUL_0443 | Phosphoenolpyruvate-protein phosphotransferase | 3/0 | 2.55 X 10<sup>-8</sup> |
| BMUL_3068 | Aldehyde dehydrogenase | 3/0 | 2.55 X 10<sup>-8</sup> |
| BMUL_4835 | Hypothetical protein | 2/1 | 2.55 X 10<sup>-8</sup> |
| BMUL_1873 | UvrD/REP helicase | 3/0 | 2.55 X 10<sup>-8</sup> |
| BMUL_2536 | Hypothetical protein | 3/0 | 2.55 X 10<sup>-8</sup> |
| BMUL_2710 | Outer membrane autotransporter | 3/0 | 2.55 X 10<sup>-8</sup> |
| BMUL_0123 | Heavy metal translocating P-type ATPase | 3/0 | 2.55 X 10<sup>-8</sup> |
| BMUL_0116 | Acyl-CoA dehydrogenase domain-containing protein | 3/0 | 2.55 X 10<sup>-8</sup> |
| BMUL_0075 | Two component transcriptional regulator | 2/1 | 2.55 X 10<sup>-8</sup> |
| BMUL_4226 | 4-hydroxyphenylpyruvate dioxygenase | 3/0 | 2.55 X 10<sup>-8</sup> |
| BMUL_4749 | Amino acid permease | 2/1 | 2.55 X 10<sup>-8</sup> |
| BMUL_4798 | Integrase catalytic region | 1/2 | 2.55 X 10<sup>-8</sup> |

<sup>a</sup> Calculated based on the probability of resampling with replacement any locus $n$ times, given a genome size of $N$ . $P = (1/N)^{(n - 1)}$ . We used $(n - 1)$ since we are calculating the probability for any locus, rather than a specific locus.
<sup>b</sup> A mutation occurred in the intergenic region flanking the start codon of this locus.
<sup>c</sup> This locus is not found in ATCC 17616 The homolog with highest similarity is in *B. cenocepacia* DDS 22E-1
<sup>d</sup> This locus is not found in ATCC 17616 The homolog with highest similarity is in *B. cenocepacia* HI2424

Some of these multi-mutated loci are known to play significant roles in antibiotic resistance. For example, a gene encoding a probable transcriptional regulator protein of MDR efflux pump cluster (BMUL_0641), which has been associated with drug resistance in multiple pathogens [31-33], has seven independently acquired mutations, and the probability of any gene being mutated seven times is 1.65×10^−23^. A locus with five multiple mutations (P = 6.48×10^−16^) encodes N-acetylmuramoyl-L-alanine amidase (AmpD, BMUL_2790), which is associated with resistance to β-lactam antibiotics [28]. Moreover, a functional enrichment analysis revealed the phosphorelay signal transduction system GO function overrepresented in multiply mutated genes compared to the functional annotation of the whole genome (P = 0.050). The phosphorelay signal transduction system has been previously described as a therapeutic target, given that it controls the expression of genes encoding virulence factors [34].

We also found ten genes that had two independent mutations located in the same or adjacent codon (Table 2). The mutational class spectrum of the SNPs associated with this observation is of 90%, 10% and 0% of non-synonymous, synonymous, and intergenic substitutions, respectively. In this case, the fraction of non-synonymous mutations is significantly higher than the fraction found for both all SNPs, as well as all the SNPs in the multiply mutated loci (P < 0.00001, chi-square test). One of the genes with multiple independent mutations in the same codon encodes for RNA polymerase sigma factor (RpoD), which is associated with the expression of housekeeping genes [35]. One of the mutations in this locus is fixed between the post-transplant isolates and the rest of the isolates, and the other mutation is fixed between the isolates in group RBG collected in the tenth sample time and the rest of the isolates.

**Table 2.** Pairs of Mutations Occurring in the Same or in Neighboring Codons.

| Encoded Protein | Proximity |
| --- | --- |
| Regulatory protein GntR, HTH:GntR, C-terminal | Adjacent codon |
| Oligopeptide ABC transporter, periplasmic oligopeptide-binding protein (OppA) | 2 codons away |
| Citrate-proton symporter | 2 codons away |
| CDP-6-deoxy-delta-3,4-glucoseen reductase-like | 2 codons away |
| RNA polymerase sigma factor (RpoD) <sup>a</sup> | Same codon |
| Endo-1,4-beta-xylanase Z precursor <sup>b</sup> | Adjacent codon |
| Isoquinoline 1-oxidoreductase beta subunit <sup>b</sup> | 2 codons away |
| LSU ribosomal protein L4p (L1e) <sup>b</sup> | Same codon |
| Chaperone protein (DnaJ) <sup>c</sup> | Adjacent codon |
| Probable transcription regulator protein of MDR efflux pump cluster <sup>d</sup> | 2 codons away |

a Loci additionally mutated 1 more time. Additional mutation is synonymous.
b Loci additionally mutated 1 more time. Additional mutation is non-synonymous.
c Locus additionally mutated 2 more times. All non-synonymous mutations.
d Locus additionally mutated 5 more times. All non-synonymous mutations.

### Parallel pathoadaptive variants are overrepresented in recombinogenic regions

We identified a minimum of 14 recombination events in our full dataset based on the four-gamete tests of Hudson and Kaplin [36] (Fig 2D). Three of these events were identified between sites in different genome assembly contigs; therefore, they were not considered in downstream recombination analysis. The nucleotide length of this recombinogenic regions ranged from 4,783 bases to 192,532 bases, and these regions account for 15.1% of the assembled genome. 300 (15.9%) out of the total 1,892 SNPs and 47 indels (14.3%) occur in these regions, which is not significantly different than expected given the recombinogenic proportion of the genome.

We next looked to see if there was an association between recombination and the evolution of antibiotic resistance. 51 (18.3%) of the 279 SNPs associated with both aminoglycosides tested (amikacin & tobramycin), and 42 (14.9%) of the 281 SNPs linked to ciprofloxacin are found in recombinogenic regions (Fig 5A). These ratios fail to reject the null hypothesis that these mutations are randomly distributed around the genome. On the other hand, 52.9% (9 of 17 SNPs) and 47.4% (9 of 19 SNPs) of the SNPs associated with aztreonam and ceftazidime, respectively, are found in recombinogenic regions, which are significantly different than expected by chance (p < 0.0001, chi square test).

**Fig 5.**
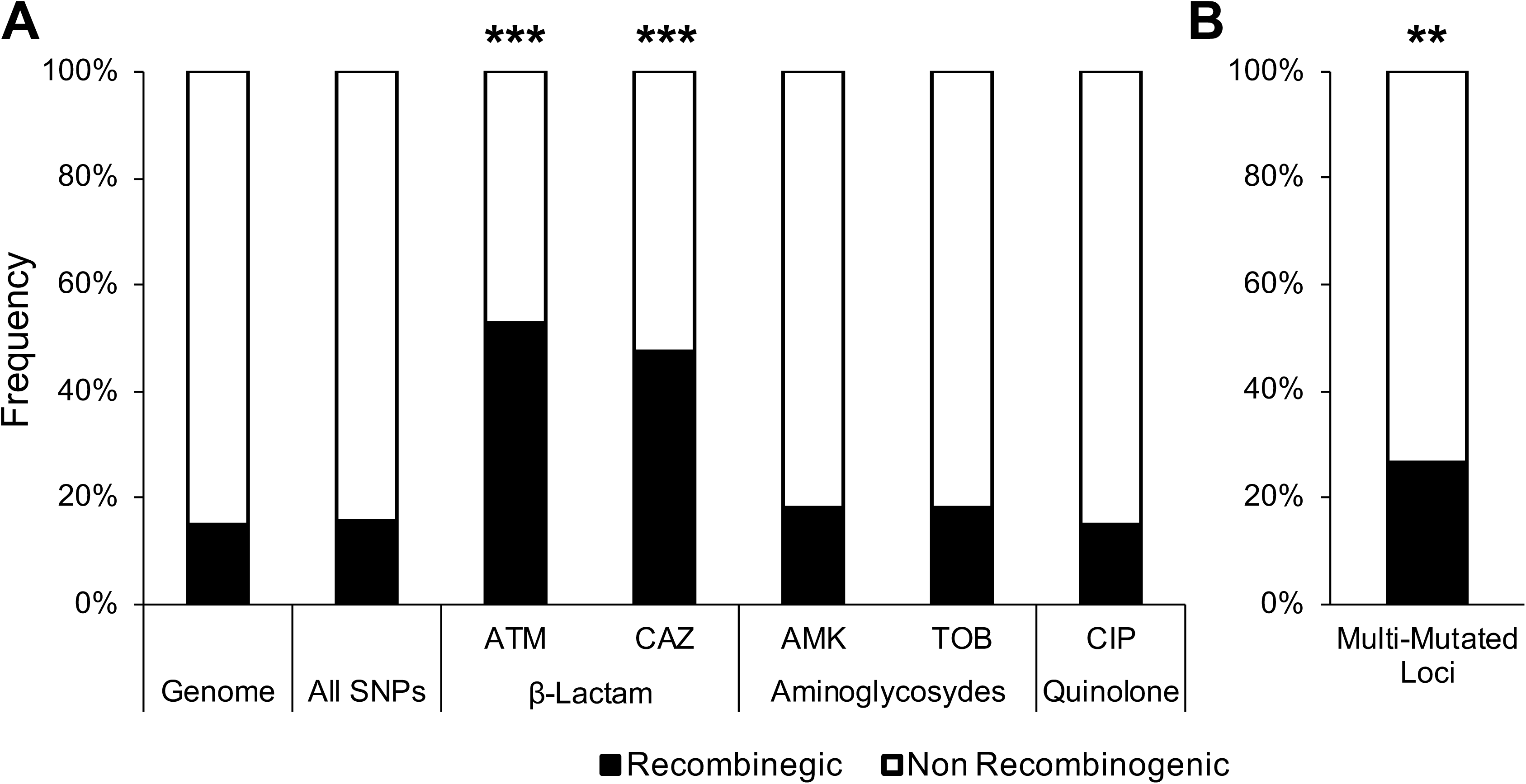
Distribution of pathoadaptive variants in recombinogenic regions of the genome. (A) Distribution of the mutations associated with the tested antibiotics in the identified recombinogenic regions and in the rest of the genome (*** p < 0.0001, chi square test with multiple test correction). (B) Distribution of the mutations in multi-mutated loci in the identified recombinogenic regions and in the rest of the genome (*** p < 0.001, chi square test with multiple test correction).

Finally, 49 (26.6%) of the 184 SNPs and 4 (8.5%) of the 47 indels found in loci independently mutated three or more times occur in the identified recombinogenic regions (Fig 5B). Thus, while SNPs involved in multi-mutated loci are overrepresented in recombinogenic regions more than expected (P < 0.0001, chi square test), indels in multi-mutated genes are not significantly underrepresented.

## Discussion

Our study investigated how *B. multivorans* evolves within the lungs of an individual afflicted with CF using a deep longitudinal sampling design (i.e. multiple isolates obtained per sputum sample) to capture both the overall population diversity and the temporal shifts that occurred at different phases of the infection, including the colonization of a new allograft. To identify the source of genetic diversity in this *B. multivorans* population, we needed to understand: 1) the genetic relationships between the incident isolate that was recovered from the first BCC-positive sputum culture, the chronic strains that persisted in the population, and the population of strains that re-established an infection post-transplant; 2) whether there were multiple colonization events of the patient by divergent clones; 3) how genetic diversity was generated and dispersed in the population; and 4) how the pathogen adapts and responds to clinical treatment. While we were unable to address all of these questions, we have concluded that the chronic population originated from either the incident isolate, or a clone that shared a recent common ancestor with the incident isolate. Furthermore, all of the chronic isolates descended from a single common ancestor, ruling out multiple independent colonization events.

One clear signal is that the *B. multivorans* isolates recovered from the post-transplant lung did not originate from the chronic population. In fact, it appears that the post-transplant isolates came from a new infection that originated from the same source as the incident infection. Unfortunately, the source of these infections cannot be determined, and could be either the environment or the patient’s upper respiratory tract. In the former case, it is likely that the patient lived in the same home or locale over the course of the study, and that the ancestral *B. multivorans* clone is endemic in that environment. Alternatively, in the latter case, the upper respiratory tract is known to act as a reservoir for a number of CF pathogens [37].

Consequently, it is possible that clonal descendants of the ancestral or incident strains resided in the patient’s upper airways since the incident infection. Some transplant procedures attempt to clean the nasal reservoir prior to transplant via nasal washing / scraping, but we do not know if this was done for this patient. If this hypothesis is true, it would explain why the post-transplant isolates have an antibiotic susceptibility pattern much more similar to the chronic isolates than the incident isolate. We also note that the post-transplant population is much more genetically diverse than any of the chronic populations. This could suggest that this population was rapidly adapting to an environmental change, such as the shift from CF to non-CF conditions, which would include, differences in immune response, the composition of the allograft microbiome, and treatment regimens. Alternatively, it could reflect colonization by a population of related strains. It is possible that given sufficient time this population would eventually be winnowed down to a single surviving clone (as is seen with the incident infection) due to selection and / or genetic drift

A major motivator for this study was to better understand how pathogens adapt to their hosts over the course of disease progression and treatment; an issue that can be addressed using statistical association tests. Correcting for the genetic structure of the bacterial population poses a challenge to the implementation of these tests. Population structure in this context refers relationships among strains due to descent form a common ancestor and limited recombination. This structure results in the linkage of segregating genetic variation around the genome, which makes it very difficult to distinguish a causal mutation that is responsible for a phenotype of interest from a neutral variant that occurred in the same genetic background. In the absence of recombination, the neutral mutation will have the same population distribution as the causal mutation due to genetic hitchhiking. This issue is particularly prevalent when studying largely isolated and recently evolved populations, such as the case of pathogens evolving within a host.

To overcome these two issues, we imposed a lineage control filter on our GWAS approach, in which we focused only on mutations that occurred in multiple, distinct, genetic lineages. This pattern can best be explained by recombination of polymorphisms between lineages, but formally, could also be due to extensive gene loss. Our analysis showed that linkage disequilibrium was only disrupted in a relatively small number of polymorphism (those polymorphisms shown as orange circles; S7b-e Fig). This reinforces the need for deep sampling since the infrequent recombination signals may have been missed if isolates were only collected from a single sample, or if only single isolates were recovered from each sample. Consequently, the tractability of GWAS in this *B. multivorans* population was greatly enhanced by our sampling schema.

Using the established lineage structure of the *B. multivorans* population as control for our association study, we identified two non-synonymous SNPs associated with resistance to the aminoglycosides amikacin and tobramycin, and to the quinolone ciprofloxacin. One of these SNPs occurs in a locus encoding the transcription factor AraC, which is involved in the global regulation of efflux pumps, while the other SNP was found in a locus annotated as a porin. Although not specific to aminoglycosides or quinolones, overexpression of efflux pumps and repression of porin proteins has been reported as important mechanisms of antibiotic resistance for bacteria [38]. Neither mutation is projected to significantly vary the function of the encoding protein.

Additionally, we identified a single SNP associated with resistance to the β-lactams aztreonam and ceftazidime. This SNP occurs in the *ampD* gene, which is a negative regulator of the β-lactamase AmpC, and it is expected to have a deleterious effect in the encoding protein. This observation is not unexpected as bacteria treated with β-lactams would benefit from the constitutive overproduction of β-lactamase. Overall, AmpD seems to play an important role in the pathoadaptation of this *B. multivorans* population since four other independent non-synonymous mutations, all of which are expected to have deleterious effects on the protein, occur at this locus (S8a Fig).

Our use of the population control criterion of only considering mutations present in multiple lineages meant that we excluded some variants associated to virulence, such as one of the four mutations in *ampD*, which was statistically associated with β-lactam resistance. Without our population control it would be impossible to identify causative mutations from hitchhiking variants that are in linkage disequilibrium with the causative mutation. Filtering in this manner reduces the number of false positives; nevertheless, variants underlying phenotypes of interest could be segregating in linkage disequilibrium blocks, and therefore, may not be identified in our GWAS approach (false negatives).

We observed that mutations associated with resistance to β-lactams (prior to lineage controls) occur disproportionately in recombinogenic regions (Fig 2F), while variants associated with both aminoglycosides or ciprofloxacin are more randomly distributed with respect to recombinogenic regions. The study patient received both long-term maintenance β-lactam and aminoglycoside treatments in addition to multiple short-term β-lactam treatments that included cycles of ceftazidime, piperacillin/tazobactam, meropenem, and cefepime. This more aggressive and varied course of treatment with β-lactams could potentially explain the increased role of recombination in the dissemination of putatively beneficial polymorphisms, similar to what has been observed in other pathogens [39, 40].

Our analysis identified genes under strong selection by focusing on loci with a statistical excess of independent mutations (i.e. parallel pathoadaptation) [25, 41, 42]. Examining multi-mutated loci can reveal the heterogeneous selective pressures that bacteria must adapt to in order to reside within the lung. For instance, a gene encoding a transcription regulator of multidrug resistance efflux pumps independently accumulated seven different mutations leading to eight unique alleles in our population of 111 *B. multivorans* isolates. We also found seven different alleles of a locus encoding cyclic β-1,2-glucan synthase, which is linked to bacteria’s ability to elude host cell defenses [43]. A number of loci underlying virulence-associated traits, such as quorum sensing and biofilm production, also carry multiple independent mutations. Particularly interesting are multiply mutated loci with no characterized function, or with no prior linkage to resistance or virulence. These loci include a NAD-glutamate dehydrogenase locus BMUL_4010, which was mutated five independent times over the course of the study, and a glycosyl transferase protein (BCEN2424_5592), not previously seen in *B. multivorans* that was mutated six times (4 SNPs and 2 indels) during the course of the study. Examples such as these provide excellent candidates for characterizing the cryptic resistome – loci previously not known to be involved in antimicrobial resistance. In addition, the strongest signals of parallel pathoadaptation involve those cases where mutations occur independently in the same or adjacent codon. These observations point to a very specific form of selective pathoadaptation, which identifies the specific residue or region of the locus that potentially plays a role in selective advantage and may affect a conserved function.

Finally, our study highlighted an intriguing role for recombination in the development of antimicrobial resistance in *B. multivorans*. We observed that multi-mutated loci were over-represented within recombinogenic regions, along with an excess of mutations associated with β-lactam resistance. This suggests that while recombination plays an important role in the pathoadaptation of this *B. multivorans* population, its selective benefit may be environment dependent.

Our study illustrates the relevance of deep, longitudinal sampling to the implementation of GWAS approaches in a population under positive selection. We identified the potential genetic basis behind the antibiotic resistance of a *B. multivorans* population in a single host. Moreover, this approach allowed us to study variants associated to antibiotic resistance and revealed that resistance to β-lactams may be passed within the population via recombination. This study is limited to *in silico* predictions of the impact mutations on protein function, and future efforts should include functional validation of these mutants; nevertheless, many of the identified genes are already well-established targets for antibiotic resistance. Additionally, our findings are restricted to a single patient and a single bacterial species; extending this approach in other systems under positive selection will be required to establish the generalizability of the findings. Nevertheless, this study is one of the first examining in depth the fine-scale evolution of *B. multivorans* in the lungs of a CF patient as it transitions from an early infection to chronic infections and the eventual reinfection of a transplanted allograft.

## Materials and Methods

### Ethics statement

All protocols involving the collection, handling and laboratory use of respiratory specimens were approved by the Research Ethics Boards of St. Michael’s Hospital (Protocol #09-289) (Toronto, Canada) and the University Health Network (Protocol #09-0420-T) (Toronto, Canada). We obtained informed consent from the study subject prior to specimen collection and sputa were produced voluntarily. All experiments involving clinical specimens were performed in accordance with the *Tri-Council Policy Statement: Ethical Conduct for Research Involving Humans*, of the Canadian Institutes of Health Research, the Natural Sciences and Engineering Research Council of Canada, and the Social Sciences and Humanities Research Council of Canada.

### Specimen collection and isolation of *B. multivorans*

Sputum specimens were collected by expectoration from a 29-year-old male (CF170), with a homozygous ΔF508 CFTR genotype being followed at the Adult CF Clinic at St. Michael’s Hospital (Toronto, Canada). Ten sputum specimens were collected over a 10-month period while the patient was in the advanced stages of CF lung disease (assessed by the forced expiratory volume in 1 second (FEV1), FEV1 which was 27-39 % predicted throughout the course of the study), and an additional sputum specimen obtained after the patient had undergone double lung transplantation. All specimens were processed for bacterial culture as previously described [44]. After 48h of incubation, cultures were visually inspected, and each distinct colony morphotype was described using eight characteristics of physical appearance (pigmentation, size, surface texture, surface sheen, opacity, mucoidy, autolysis and margin shape). Ten colonies were selected from each sputum culture in relation to the diversity of colony types present. The incident isolate was obtained from the *Burkholderia cepacia* complex repository at St. Michael’s Hospital and was recovered from the first BCC positive sputum culture produced by the study patient (Toronto, Canada). Isolates were stored at −80°C in 20% (v/v) glycerol after a 20h subculture in LB broth (Wisent Inc., QC, CA) and confirmed as *Burkholderia* spp. by a secondary subculture onto both *Burkholderia cepacia* selective (BCSA) (HiMedia Laboratories, Mumbai, IN) and MacConkey (Becton Dickinson, MD, USA) agars, as well as being tested for growth at 42°C. The *recA* gene was sequenced from each isolate as described by Spilker *et al*. for preliminary speciation [45].

### Antimicrobial susceptibility testing

Each isolate confirmed as *B. multivorans* was screened for antimicrobial susceptibility by agar dilution using Clinical and Laboratory Standards Institute procedures [46]. We tested susceptibility to representatives of the β-lactam (aztreonam [ATM], ceftazidime [CAZ]), fluoroquinolone (ciprofloxacin [CIP]) and aminoglycoside (amikacin [AMK], tobramycin [TOB]) (Sigma-Aldrich, ON, Canada) classes. Minimum inhibitory concentrations (MIC), defined as the lowest concentration of each antibiotic to inhibit growth, were reported as the median MIC of three independent experiments. Growth was assessed following 24 to 48 h of incubation on Mueller-Hinton agar (Becton, Dickinson, MD, USA). The *B. multivorans* ATCC 17616 strain was included as a positive control, while *P. aeruginosa* ATCC 27853 and *E. coli* ATCC 25922 were used as quality controls.

### Sequencing and Quality Control

*B. multivorans* isolates were whole-genome sequenced on the MiSeq and NextSeq Illumina platforms. The number of bases sequenced per isolate ranged from 213 to 2,262 million bases, and the median was 1,002 million bases. Trimmomatic v. 0.33 was used to remove adapters and quality trim the sequencing reads from each isolate (parameter settings: PE -phred33 ILLUMINACLIP:adapters.fa:2:30:10 LEADING:5 TRAILING:5 SLIDINGWINDOW:4:25) [47]. Sequencing reads with guanine homopolymers longer than ten bases were trimmed with cutadapt v. 1.9.1 (parameter settings: −a “G{10}”) [48]. Reads bellow 100 bases were removed using Trimmomatic v. 0.33 (parameter settings: PE −phred33 MINLENGTH:100). The resulting quality-controlled sequencing reads yielded a median read depth per position of 117X (range 32-276X).

### *De novo* and Reference Mapping Assembly

Each of the isolates was *de novo* assembled using the CLC Genomics Workbench v. 8.0.1 (Aarhus, Denmark). Contigs with a scaffolding depth lower than 10X and/or with a size smaller than 1 Kb were removed from further analyses. Isolate CF170-3b, which was sequenced with 250 bp-long paired-end reads, yielded the best assembly metrics in 26 contigs with lengths ranging from 1,010 to 1,243,078 bases and an N50 of 654,231. The final assembly length of the CF170-3b isolate was of 6,444,123 bp. These contigs were annotated at the RAST server using the native gene caller and Classic RAST as the annotation scheme [47]. Further, this genome was functionally annotated with blast2go v 4.1.9 [49] including blastx v. 2.6.0+ [50]. Statistical results from the functional enrichment analysis were Bonferroni corrected for multiple testing using the number of multiply-mutated genes (P-value/62). The contigs of the CF170-3b genome were used as the reference for mapping assembly of each remaining isolate. We performed three different reference-mapping assemblies including BWA v 0.7.12 [51], LAST v 284v [52] and novoalign v 2.08.03 (Novocraft Technologies).

### Single Nucleotide Polymorphism (SNP) and indel Calling

SAMtools and BCFtools v 0.1.19 were used to produce the initial set of variants [53]. We implemented a method previously described to detect SNPs among the 111 isolates [25, 54]. First, 1,892 high-confidence polymorphic positions were identified using the following criteria: 1) variant Phred quality score of ≥ 30 and 2) variants must be found at least 150 bp away from either the edge of the reference contig or an indel. Second, we reviewed each high-confidence polymorphic position in each isolate with a relaxed Phred score threshold of 25. Support for either the reference or the SNP call was verified with a multi-hypothesis correction which required that at least 80% of the sequencing reads endorsed the SNP or the reference. If the data did not support either base, then the position was called as an ambiguous base (‘N’). The ambiguous call rate was lower than 0.01%.

Candidate indels detected by BWA and SAMtools were examined by realigning mapped and unmapped sequencing reads to the indel regions using Dindel v. 1.01 [55]. High-confidence indel positions were defined as sites with: 1) variant Phred quality score of ≥ 35; 2) at least two forward and two reverse reads; and 3) sequencing coverage ≥ 10. These indel positions were reviewed in each isolate. The final indel call required a Phred quality score ≥ 25 and an allele frequency ≥ 80%. Ambiguous indel calls were defined as those where the allele frequency was ≤ 20%.

### Population and Single Genome Sequencing Evaluation

We performed bulk population sequencing on the post-transplant specimen to confirm that our isolate sampling depth appropriately represented the real *B. multivorans* population diversity (S9 Fig). The sequencing reads from each of the ten isolates from the post-transplant sample were rarified to 1/10^th^ of the number of sequencing reads produced by the population sequencing experiment. These reads were combined in corresponding paired-end fasta files. Next, population and single isolate sequencing reads were mapped to the *de novo* assembled genome of the CF170-3b isolate using BWA. Mutation allele frequencies for each experiment were estimated as previously described by Lieberman *et al*. [54].

### Phylogenetic, Population Structure, Coalescent and Recombination Analyses

Using the 1,892 SNPs, we created a genome-wide alignment to reconstruct the phylogenetic relationships among the 111 isolates. The phylogeny was calculated using MrBayes v. 3.2.6 [56]. The nucleotide substitution model that best fit our data was the General Time Reversible (GTR) with gamma-distributed rate variation across sites (LnL=-13,152.7810, AIC= 26,832.1306) as calculated with jModelTest v. 2.1.10 [57]. The Bayesian analysis was run through four different chains of 1 million Markov Chain Monte Carlo (MCMC) generations sampled every 100 MCMC generations and the burn-in period was of 250,000 MCMC generations. The final average standard deviation of split frequencies was of 7.3×10^−3^, and the potential scale reduction factor (PSRF) of the substitution model parameters ranged from 1 − 6.66×10^−5^ to 1 + 4.83×10^−4^. The phylogeny was rooted with *B. multivorans* ATCC 17616 [58]. The network-based phylogenetic analysis was performed using SplitsTree v 4.14.4 [59]. We employed the Jukes-Cantor distance matrix to implement the neighbor-net Network (Fit=99.804).

The variance among the 111 isolates, including SNPs and indels, was employed to investigate the population structure using the Structure software v 2.3.4 [60]. Structure employs a Bayesian algorithm to detect the number of ancestral populations (K), also known as clusters, which describe the variance and covariance observed in a test population. The number of clusters ranging from 1-10 was tested in triplicates through 1 million MCMC generations sampled every 1,000 MCMC generations and a burn-in period of 250,000 MCMC generations. We used the correlated allele frequencies model, and admixture was allowed in these analyses. We plotted the estimated ln probability of data for the tested levels of K, and identified the smallest stable K as the optimum value since it maximized the global likelihood of the data (S10 Fig) [61]. The estimated ln probability of data plateaus at K=3, where the variance of ln likelihood ranges from 2,343.0 to 2,353.1. Assuming three ancestral populations, the isolates were classified into five different groups according to their ancestry. Isolates whose ancestry is attributed exclusively (>90%) to either ancestral population one, two, or three are grouped in group red (R), (B), or (G), respectively. Group RB includes isolates with admixed ancestry from clusters one and two (at least 10% of both cluster one and two, and less than 10% of cluster three). Isolates whose ancestral composition is made up from a combination of all three clusters (at least 10% of each cluster) are in group RBG.

We used BEAST v. 1.8.4 to implement a Bayesian approach to inferring the time to the most recent common ancestor (tMRCA) for the entire population and each group individually [62]. Next, we employed the GTR nucleotide substitution model, and estimated the nucleotide substitution frequencies with MEGA7 using the Maximum Likelihood Estimate of the Substitution Matrix tool ([AC] = 0.0091, [AG] = 0.4281, [AT] = 0.0016, [CG] = 0.0260, [GT] = 0.0061, and [CT] = 0.5290). Preliminary analyses consisting of duplicate 10 million generations and a 10% burn-in were used to estimate the appropriate molecular clock and demographic models. We tested the Bayesian skygrid, constant size and the exponential, logarithmic and expansion growth population size models using three different molecular clock models (strict and the lognormal and exponential uncorrelated relaxed clocks). The exponential relaxed uncorrelated molecular clock and the Bayesian skygrid model was inferred the most appropriate given our data ([AIC] = 26,228.421) [63]. The final analysis was run in duplicate for 1 billion MCMC generations sampled every 1,000 MCMC generation, and the burn-in period was set at 20% of the MCMC generations.

Population genetic tests and detection of recombination events in each contig were performed with DnaSP v. 5.10.01 [64].

### SNP to Phenotype Association

We tested the null hypothesis that the presence or absence of each of the 1,892 SNPs, summarized in 150 distinct mutational profiles, is equally likely found in antibiotic resistant isolates using Fisher’s exact test. These tests were conducted for each examined antibiotic at six different MIC resistance thresholds (≤16, 32, 64, 128, 256 and ≤512 MIC). For each test, we created a contingency table reflecting the distribution of each mutation profile in isolates with lower and greater MIC than each resistance threshold. *P* values were adjusted based on the total number of tests (number of mutational profiles), and only associations with a *P* value < 3.36 × 10^−4^ (0.05 / 150) were considered significant to control for multiple testing. Next, we simulated gains or losses of these mutational events following a continuous-time Markov chain along a ClonalFrameML v. 1.0-19 phylogeny as implemented in GLOOME v. 01.266 using the default parameters [65, 66]. We defined independent mutational events as those with a probability greater than 0.95 and to control for population structure, we required multiple independent mutational events in at least two STRUCTURE-defined groups.

### *In silico* mutation impact prediction

To predict the potential impact of non-synonymous SNPs on the biological function of a protein, we employed PROVEAN v. 1.1.3 [67]. These calculations were performed on the GPC supercomputer at the SciNet HPC Consortium [68].

## Acknowledgements

This research was funded by an Emerging Team Grant awarded to D.S.G from the Canadian Institutes of Health Research (CIHR) and Cystic Fibrosis Canada (CMF108027). J.D.C. was supported by an Ontario Trillium Scholarship. S.T.C. was supported by an Ontario Graduate Scholarship. B.C. was supported by a CIHR Fellowship. We would like to thank Dr. Tami Lieberman for her assistance with the estimation of the dN/dS rates.

## Supporting Information

**S1 Fig. Sequencing coverage.** Whole genome sequencing of 111 isolates of *B. multivorans* in the Illumina platform. (A) Distribution of number of bases sequenced per isolate. (B) Distribution of median read depth per position.

**S2 Fig. Genetic diversity over time.** (A) Pairwise nucleotide differences between isolates collected from the same collection sample. Incident infection is not included since only one isolate was recovered from that time point. (B) Nucleotide differences between each isolate and the incident infection isolate.

**S3 Fig. Neighbor-Net phylogeny.** This network-based phylogeny was calculated in SplitsTree v. 4.14.4. Individual strain names at the tips of each branch have been replaced with pie charts indicating the distribution of dates during which the strains were sampled (indicated by the circular legend).

**S4 Fig. Genetic diversity and selection analysis per group.** (A) Pairwise nucleotide differences between isolates from the same group based on ancestry. (B) dN/dS per group calculated including all SNPs and using only SNPs observed in multiple time points (MTP). dN/dS and the respective confidence intervals were calculated as described by Lieberman *et al*. [72]. (C). Time to Most Recent Common Ancestry (tMRCA) as estimated using the BEAST software for each group. The x axis represents the log of the years before the last sampling time. The whiskers for each data point show the 95% high probability density intervals.

**S5 Fig. SNP positions with identical distribution of reference or alternative bases across the strain collection are grouped into mutational profiles.** Here, “0”s and “1”s represent the reference or alternative base, respectively, at each SNP position for each strain. SNP1 is the only position where only Strain1 has a base alternative to the reference. Hence, mutational profile 1, 1–0–0–0, comprises only one SNP. On the other hand, Strain4 is the only strain with a variant base for positions SNP2 and SNP3. Therefore, mutational profile 2, 0–0–0–1, comprises SNP2 and SNP3.

**S6 Fig. Mutational profiles associated with antibiotic resistance.** (A) Maximum Likelihood phylogeny of 111 *B. multivorans* isolates was elaborated using RaxML v. 7.0.4 with a GTR + gamma model and 1,000 bootstraps [73]. Here, we show all mutation profiles associated with antibiotic resistance prior to lineage control in black and with lineage control in orange. (B) resistance to both β-lactams, (C) to amikacin only, (D) to both aminoglycosides, (E) to both aminoglycosides and to ciprofloxacin, (F) and to ciprofloxacin only. A filled circle represents a SNP call in the corresponding isolate compared to the reference.

**S7 Fig. Resistance levels at which genetic associations are statistically significant.** Mutational profiles were tested for association against six levels of antibiotic resistance (<16, <32, <64, <128, <256 and <512 MIC) to five antibiotics (amikacin, tobramycin, aztreonam, ceftazidime and ciprofloxacin). Black boxes show the levels of resistance at which the mutational profiles were statistically significant including multi-testing correction. Associations to ciprofloxacin antibiotic resistance are shown up to <128 MIC since no isolate had a MIC of 256 or greater in relation to that antibiotic.

**S8 Fig. Mutations in *ampD* locus.** (A) Distribution of the PROVEAN scores of all identified non-synonymous substitutions highlighting SNPs in multi-mutated loci (yellow) and in the *ampD* gene (red or blue if associated to β -lactam resistance). Red lines represent thresholds from most specific (highest), to most sensitive (lowest) to determine if a mutation is deleterious to the function of the gene in which it occurs. (B) Crystal structure of protein product of AmpD (PDB ID:2Y2B, [74]) in complex with reaction products. Mutations found in our *B. multivorans* population are colored in red or blue (mutations associated with β-lactam resistance).

**S9 Fig. Population and single isolate sequencing.** Sequencing reads from each isolate from the post-transplant sample were rarified to 1/10th of the number of reads in the population sequencing experiment; then they were combined so that the number of reads would be the same for both experiments. Sequencing reads from the population and single isolate experiments were mapped to the same reference as described above. Mutation allele frequencies for both experiments were calculated using the quality thresholds described by Lieberman *et al*. [54]. (A) Grey circles represent mutation allele frequencies in the deep population sequencing experiment (y axis) versus in single isolate sequencing (x axis). The dashed line represents the *x=y* function and the solid line is the best fit line taking into account all data points (R^2^=0.9928, 95% confidence interval= 0.9918–0.9937). Red circles represent alleles found in the single isolate sequencing experiment but not in the deep sequencing one. Fixed mutations between the reference and all the post-transplant isolates are colored blue. (B) Proportion of false positives in the single isolate sequencing experiment.

**S10 Fig. Determining the number of ancestral populations that explain the variance and covariance in CF170 *B. multivorans* population.** (A) We ran three independent chains for each K between one and ten. The estimated ln probability of data plateaus at K=3 in all chains.

